# Evaluation of *Trichoderma harzianum* mutant lines in the resistance induction against white mold and growth promotion of common bean

**DOI:** 10.1101/713776

**Authors:** Renata Silva Brandão, Thiago Fernandes Qualhato, Paula Arielle Mendes Ribeiro Valdisser, Marcio Vinicius de C. B. Côrtes, Pabline Marinho Vieira, Roberto Nascimento Silva, Murillo Lobo Junior, Cirano José Ulhoa

## Abstract

Genetic engineering has brought improvements of *Trichoderma* species for biotechnological processes such as agriculture. It has previously been reported that *sm1* and *aquaglyceroporin* genes from *Trichoderma harzianum* are strongly expressed during pathogens biocontrol. We have previously shown that *Sm1* plays a significant role in plant disease resistance and aquaporin (AQP) regulate physiological processes and responses related to water stress. Here, we report the evaluation of mutant lines with *sm1* (deleated) and *aqp* (overexpressed) in *Phaseoulus vulgaris* growth promotion and disease resistance induction. It was investigated plants physiological and biochemical responses, plant growth promotion and biocontrol of *Sclerotinia sclerotiorum*, the causal agent of white mold. Treatments included *Trichoderma harzianum* wyld type, mutant line overexpressing aquaporin (*Aqua)*, and deleted *ΔEpl1*mutant line. Our results demonstrated that Aqua mutant line increased leaf area in 25%, in comparison to non-treated plants. It also differed from other treatments, in comparison to plants with treatments ALL-42 and *ΔEpl1*, which increased 28% and 91%, respectively (Isso é abstract, ta confuse e extensor. Specific activity of β-1.3 glucanase was higher in plants treated with *T. harzianum ΔEpl1* mutant isolate, in leaves and roots respectively with 2.07 Umg^−1^ and 2.57 Umg^−1^. Plants inoculated or not with *S. sclerotiorum* presented with 2.27 Umg^-1^ and 2.34 Umg^-1^ of β-1.3 glucanase on average, respectively, but enzymatic activity was higher on leaves when compared to the roots. The *Aqua* mutant demonstrated excellent growth promoting properties. Both mutants of *T. harzianum* do not induce resistance to white mold on common bean plants.

## 1. Introduction

The common bean (*Phaseolus vulgaris* L.) is a staple food crop with an important role in human nutrition, especially as a protein source in developing countries. Currently, Brazil is the second largest producer of beans in the world, producing approximately three million tons per year, surpassed only by India. However, white mold is one of the major diseases that affect common beans and is caused by *Sclerotinia sclerotiorum* (Lib.) De Bary, a worldwide-distributed soil-borne fungus that attacks over than 408 hosts (LEITE *et al.*, 2000). Many cropped plants, such as common beans, soybean, cotton, lettuce, cabbage, tomato, sunflower, peanut and pea are susceptible to this pathogen. This pathogen is especially harmful to host plants because of lack of resistant genotypes and long-lasting survival in soil, by means of resistance structures named sclerotia. It is introduced into new areas mainly through seeds infected with the pathogen mycelium or sclerotia mixed with seeds (LIMA et al., 1998).

Sclerotia of *S. sclerotiorum* survive in soil for several years, and successful white mold management demands the reduction of the initial inoculum. Biocontrol with species of *Trichoderma* spp. have been successfully used to manage the disease, once they attack the resistance structures in soil and protect plants through different modes of action, such as hyperparastism, antibiosis and competition. (HARMAN et al. 2004; WOO et al., 2006). In this way, some *Trichoderma* isolates have been referred as plant growth promoters, according to their ability to solubilize phosphate and making them available to plants, and by the production of giberelin and auxin analogues. Such substances induce cell multiplication and elongation in plants. Further, *Trichoderma* spp. may release siderophore compounds – resulting in the solubilization of the iron present in the soil, for the benefit of plants (VINALE et al., 2008).

More recently, induction of plant resistance by bioagents has been addressed as an important mechanism to manage plant diseases, and an open field to investigate newer possibilities of *Trichoderma spp*. on plant protection. This phenomenon, called systemic resistance induction (SRI) or acquired systemic resistance (ASR) implicates in the activation of plant defense mechanisms, in response to exposure to biotic or abiotic agents. These resistance mechanisms may include the accumulation of phytoalexins and proteins related to the pathogenesis process, such as β1.3 glucanases, chitinases and peroxidase (KUHN & PASCHOLATI et al., 2010).

There are several possibilities to improve SRI, ARS and biocontrol gains, with the exploration of antagonist diversity, formulation methods, and methods of delivering bioagents in the field. Antagonist diversity can be achieved with traditional methods of antagonist selection, or even with breeding methods that include protoplast fusion and chemically driven mutations. *Trichoderma* mutant isolates that either overexpress specific proteins or have genomic deletions may be not only recommended for protection against biotic threats, but also assist plants on their protection against abiotic hazards, such as osmotic shock (TANGHE et al., 2006).

For instance, a deletion of the gene Δ*Epl1/Sm1* in *Trichoderma harzianum* resulted in loss of biocontrol capacity against phytopathogenic fungi, with the mutant loosing capacity of recognition, coiling in host hyphae and expression of defense related to bean plants. The *ΔEpl1* gene expresses cerato-platanin proteins released during the early stages of filamentous fungi development (BODDI et al., 2004; PAZZAGLI et al., 1999; SCALA et al., 2004). However, there are no results concerning the consequences of this gene deletion on the induction defense responses in plants, once Sm1 or *ΔEpl1* proteins, respectively from *T. virens* and *T. atroviride* induce phytoalexin production and/or cell death in host and non-host plants (BODDI *et al*., 2004). Aquaporins are water channel proteins that increase the permeability of the cell membrane lipid bilayer to water. Although the movement of water through the cell membrane occurs directly through the lipid bilayer (simple diffusion), in some cells most of the osmosis is facilitated by these integrated proteins, the aquaporins (facilitated diffusion), these proteins contain a simple, selective pore to water, which allows the rapid passage of this molecule through the membrane by facilitated diffusion. Each aquaporin allows the entry of 3×109 molecules of water per second. Without these proteins, only a small fraction of these water molecules would diffuse through the same area of the cell membrane in a second (CAMPBELL, 2008).

Biochemical interactions between *Trichoderma* transformants and pathogens have being the subject of several reports, as the increasing availability of *Trichoderma* species genomes have allowed the functional analysis of genes involved in both biocontrol and defense elicitors in plants, in a larger scale (VIEIRA *et al.*, 2017). Such biochemical investigation is necessary to compare biocontrol capacity of mutants with their wild strain parents, to detect morphological alterations and to measure enzymatic activities that explain phenotypical differences (GRUBER *et al.*, 2012). Up today, changes in growth promotion and protection against diseases were not investigated regarding common bean plants and *Trichoderma* mutants. Therefore, the objective of this study was to evaluate the physiological, anatomical and biochemical effects of the fungus *Trichoderma* mutant *Aqua* and *ΔEpl1* in common bean plants against *S. sclerotiorum*.

## 2. Materials & methods

The experiments were carried out at Enzimology Laboratory of Universidade Federal de Goiás (Goiânia, Brazil) and at Embrapa Arroz e Feijão (Santo Antônio de Goiás, Brazil). The tests were performed with three *Trichoderma harzianum* isolates: the ALL 42 wild strain and their mutated strains “*Aqua*” (which overexpresses the aquaporin protein) and *“ΔEpl*” (this gene is involved in the process of mycoparasitism against phytopathogenic fungi).

### 2.1. Inoculum production

The *Trichoderma* strains were cultivated on 250 ml Erlenmeyer flasks, with 100g of parboiled rice grains previously moistened with 60% distilled water and autoclaved (121 °C, 40 min). Flasks were incubated for seven days at 25 °C and 12-hour photoperiod. After that, conidia were harvested with the addition of distilled water and filtered with a cheesecloth, to adjust spore suspensions at 2 × 10^9^ conidia mL^-1^, with the help of a hemocytometer.

*Trichoderma* conidial suspensions were employed to microbiolize seeds of ‘BRS Estilo’, a popular common bean cultivar in Brazil, with carioca type grains. The seed treatments consisted on seed immersion in spore suspensions for 10 minutes, followed by a two-hour drying period at room temperature. Control treatment seeds were not inoculated. Afterwards, a greenhouse experiment was set up with treatments sown in 5 kg pots with soil amended with a balanced nutrient solution, to meet the common bean nutritional requirements. The pots were watered daily, and additional fertilization of plants with the same nutrient solution was carried out weekly. The internal greenhouse maximum temperature was adjusted to 25°C with the help of a thermostat, which started a cooling system (fan and moisturized expanded clay) when necessary. The experimental design was completely random, with three replicates for each treatment. The plants were grown free of pests and other pathogens and harvested at R6 stage, corresponding to the blooming period, 47 days after sowing.

The plants were inoculated with the pathogen with BDA disks, 6 mm in diameter, to the leaflet and the junction of the third trifolium, and the discs were deposited so that the mycelium was in contact with the plant. The culture medium was drilled with the very tip used for inoculation, which was completely filled with the disks. Subsequently, the plants were kept in a greenhouse for 8 days at 25 ± 5 °C and at a relative humidity of more than 85%.

The whole plants (aerial part and the root system) were harvested, with their roots thoroughly washed in tap water to remove soil articles, with five plants immediately sent for physiological analysis. Five leaves and roots of each plant were kept and soon after stored in a -80 °C freezer, for enzymatic analysis.

### 2.2. Physiological analysis of leaf area, root system and dry mass

The leaf area of common bean plants was estimated with a LI3100 (LiCor) equipment, and the root system morphology was assessed with WinRHIZO Pro 2007 (Regent Instruments. Inc.) software, coupled with an Epson XL 10000 professional scanner equipped with an additional lighting unit to improve image quality. Plant roots were carefully distributed in a transparent acrylic tray (20 cm × 30 cm) filled with an approximately 1 cm water layer, supporting image acquisition and measures of root length, diameter, surface and volume. Finally, plant dry mass was estimated with samples dried at 60 °C, for approximately 72 hours.

### 2.3. Enzymatic Assays

The common bean leaves previously stored at -80 °C were macerated in liquid nitrogen with the help of pistil and a mortar, until achieving a fine powder. The sample transferred to a 1.5 ml microtube with a 1:4 (v/v) extraction solution composed of Tris-HCl 10 mM; NaCl 150 mM; EDTA 2 mM; DTT 2 mM; PMSF 1 mM; Leptina 10 mg.ml-1 e Apotinina 10 mg.ml-1. The suspension formed was stirred for five minutes and centrifuged under refrigeration for 30 minutes at 13,000 × g. The supernatant was discharded, for the following enzymatic tests.

#### 2.3.1. β-1.3 glucanase

Activity of β-1.3 glucanase enzyme was measured in a reaction medium formed by 50 μL of crude plant extract and 1% laminarin in 1.0 M sodium acetate buffer (pH 4.5), which was incubated at 35 °C. Initial reaction speed were studied. Quantification of reaction products was realized using the DNS (3.5-dinitrosalicylic acid) 1% method, with the aid of a FEMTO 600 Plus spectrophotometer, at 540 nm. A standard glucose curve was obtained. Specific enzymatic activity was estimated as the amount of glucose (in mg) produced in reaction medium per time per minute and per milligram of protein (mg of glucose.min^−1^.mg^−1^). One milligram of glucose produced per minute was defined as a unit (1U) of enzymatic activity.

#### 2.3.2 Peroxidase

Peroxidase enzyme activity was carried out in a reaction medium composed of 10 μL of crude plant extract, 0.5% hydrogen peroxide and 2,2’-azino-bis (3-ethylbenzthiazoline-6-sulfonic acid) (ABTS) in 1.0 M sodium acetate buffer (pH 4.5), incubated at 30 °C, where initial reaction speed was observed. Quantification of reaction product was carried out in Spectrum SP-2000UV spectrophotometer, at 405 nm. Concentrations of the reaction product were estimated using molar extinction coefficient of the product (3.6 × 10^4^ mol^−1^.cm^−1^). Specific enzymatic activity was defined as the amount of product in μmol produced in reaction medium in seconds and per milligram of protein (μmol of ABTS * .s^-1^.mg^-1^). One μmol of ABTS produced per second was defined as a unit (1U) of enzymatic activity.

#### 2.3.3. Chitinase

Chitinase enzymatic activity was also performed in a reaction medium composed of 50 μL crude extract of plant leaves, 0.25% chitin solution 1.0 mM, in 1.0 M sodium acetate buffer (pH 4.5), incubated at 35 °C. Initial reaction speed was analyzed by quantification of the reaction product, which was performed using the DNS (3.5-dinitrosalicyclic acid) 1% method. Results were observed with FEMTO 600 Plus spectrophotometer, at 540 nm. A standard glucose curve was drawn, and specific enzymatic activity was defined as the amount of glucose in mg produced in reaction medium in minutes and per milligram of protein (mg of glucose.min^-1^.mg-1). One milligram of reducing sugar produced per minute was defined as a unit (1U) of enzymatic activity.

#### 2.3.4. Phenylalanine ammonia-lyase

Phenylalanine ammonia-lyase activity was performed with a crude extract of plant leaves homogenized in sodium borate buffer (pH 8.8) and in 20 mM L-phenylalanine. Trans-cinnamic acid derivatives absorbance was measured in a spectrophotometer at 290 nm.

#### 2.3.5. Lipoxygenase

Lipoxygenase activity was measured by a spectrophotometer using 10 mM linoleic acid as substrate, according to the methodology reported by Rangel et al. (2002). Increase of absorbance at 234 nm, resulting from the formation of a conjugated double bond system in the hydroperoxide produced by the reaction, was quantified. One unit of lipoxygenase activity was defined as the amount of enzyme generating 1 μmol of hydroperoxides per minute.

### 2.4. Statistical analysis

Experimental data was firstly subjected to univariate analysis of variance, with each variable appraised individually. When significant differences of 5% were detected, the Duncan test (5%) was performed, to separate the averages of treatments. In sequence, the complete data set was subjected to principal component analysis (PCA) with PAST 3.0 software (University of Oslo, Norway), to verify the contribution of each variable on the whole experimental variance, and estimate the relationships between plant development and enzyme production, affected by the treatments.

## 3. Results and discussion

### 3.1 Physiological results

The *Aqua* treatment promoted a 25% increase in the leaf area of plants compared to the control, differing from other treatments. Plants from seeds treated with ALL-42 and *ΔEpl1* isolates showed 28% and 91% increase, respectively. Control plants presented 2.3% higher leaf area than plants exposed to ALL-42 treatment and 54% higher than plants with *ΔEpl1* treatment. (figure 1)

**Figure 1:**
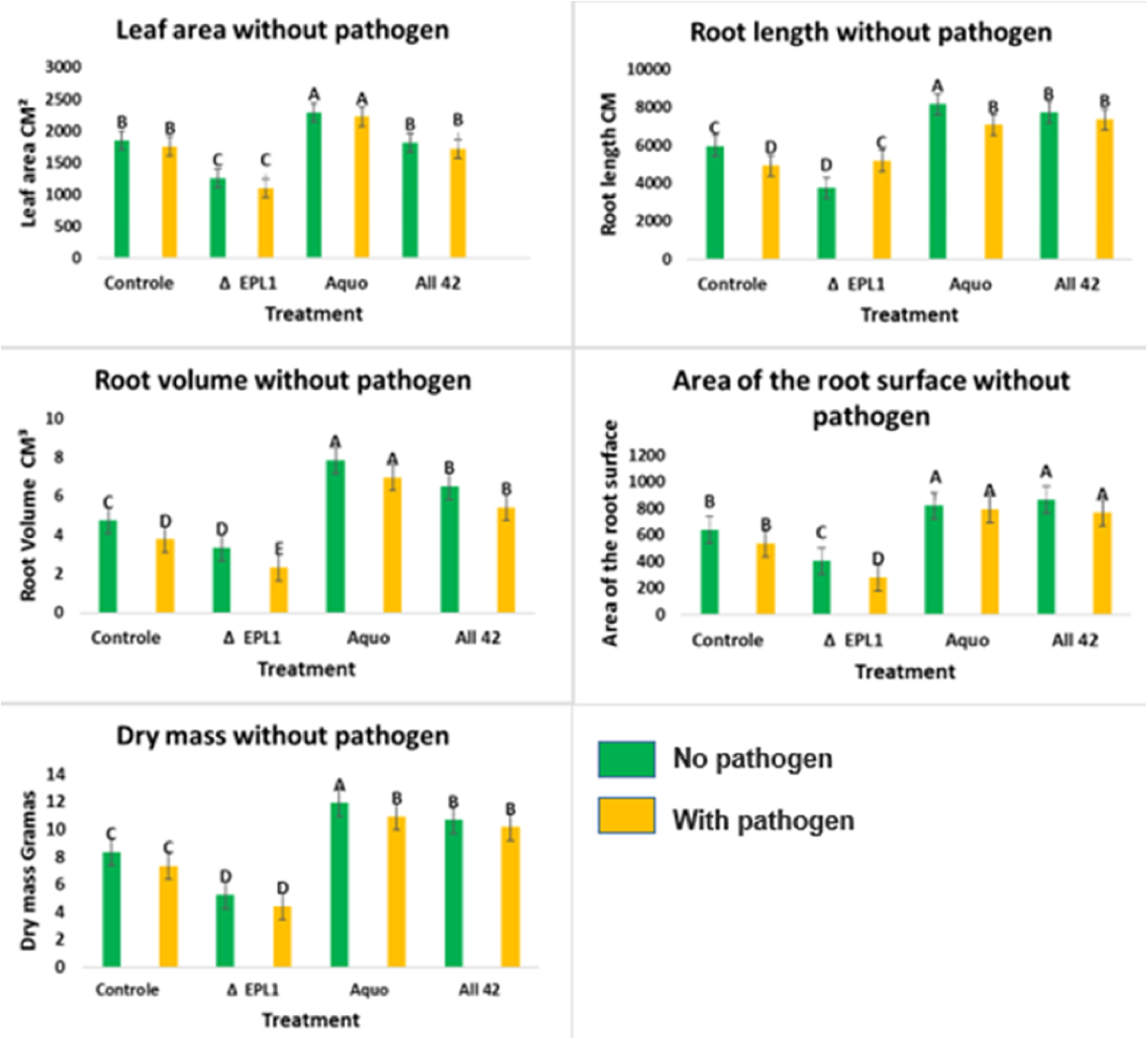
Effect of *Trichoderma harzianum* isolates on leaf area, root length, root volume, surface area and dry mass in ‘BRS Estilo”common bean plants with and without the pathogen *Sclerotinia sclerotiorum.* Averages of the same, do not differ among them by the test Duncan to 5% of significance.

Root length was 40% higher in relation to control for plants from the *Aqua* and ALL-42 treatments, without differences between them. In plants originated from *ΔEpl1* treatment, root length was 65% smaller when compared to control and 131% smaller in relation to the other treatments. Root volume of plants with the *Aqua* treatment was 25%, 75% and 169% higher, when compared to plants assigned with ALL-42, control and *ΔEpl1* treatments, respectively. Plants in the ALL-42 treatment had a 40% larger root volume and plants with *ΔEpl1* treatment presented 67% lower root volume when compared to control plants. (figure 1)

When compared, plants related to ALL-42 isolate and *Aqua* mutant strain treatments did not differ in relation to root surface (815.63 cm^2^ and 807.21 cm^2^, respectively). They showed a 39% increase in root surface of plants with ALL-42 treatment, and 37% for the *Aqua* mutant lineage treatment, when compared to control. Root surface for plants treated with *ΔEpl1* mutant isolate were 72% lower than control plants. (figure 1)

Regarding plant dry matter, the highest gains were achieved with plants treated with *Aqua* mutant lineage and with isolate ALL-42, with increases of 11.42 g and 10.43g, respectively. Plants treated with mutant isolate *ΔEpl1* led to lower results, of 4.83 g in contrast to control plants with 7.83g on average. (figure 1) Pereira et al. (2014) reported the interactions between *T. harzianum* and common bean grown in soil infested or not by the root pathogens *F. solani* and *R. solani*. This study showed that the antagonist could promote growth of bean plants, increasing plant height, leaf and radicular area, along with the number of secondary roots, with shifts on its root system architecture. Such changes are favored by several mechanisms that modulate plant growth and development, which in combination with plant genotype can result in different plant phenotypes, indicating that *Trichoderma* genotype is also important for modulating plant phenotype (SALAS-MARINA et at., 2015). Several reports showed that *Trichoderma* beneficial effects on plants occurs through the production of phytohormones, phytohormone-like molecules, volatile organic compounds, secondary metabolites or plant phytohormone homeostasis (SALAS-MARINA et al., 2015, OLMEDO-MONFIL and CASAS-FLORES, 2014, SÁENZ-MATA et al., 2014).

Growth is promoted by microorganisms and occurs through the action of several unclear factors, involving phytohormones and vitamins production or material conversion into a form for the plant to use, minerals absorption and transport and pathogens control. Several studies have shown the beneficial effect of *Trichoderma* species on plant development.

Carvalho Filho et al. 2008, concluded that *Trichoderma spp* CEN 262 isolate provided a higher development index in eucalyptus seedlings. Studies with *Arabidopsis thaliana* investigated the role of auxin produced and isolated from *Trichoderma spp*. in plant growth and development in response to *T. virens* and *T. atroviride* inoculation, leading to a fungus-plant interaction system, resulting in phenotypic characteristics related to auxin, such as increased biomass production and stimulation of root development (CONTRERAS-CORNEJO, 2009).

Carvalho et al. 2011, conducted an experiment with six *Trichoderma spp.* isolates, in the promotion of bean plant growth. The group observed that four of the isolates increased dry matter of plants aerial parts, ranging from 4.42 to 5.71%. Some of the *Trichoderma* isolates used in this work also provided increase of the plants. Brotman et al. 2010, also used *Trichoderma* species that promoted increases up to 300% in plant growth. Beneficial effects of these fungi had been reported in several important crops development, such as papaya, tomato, soybean, corn, cucumber, peppers and eucalyptus (CARVALHO FILHO et al., 2008, FONTENELLE et al. SILVA et al., 2011).

Chagas et al., 2017, worked with different crops such as maize, rice, soybean and cowpea, and *T. asperellum UFT 201* inoculation was superior to biomass characteristics, revealing its potential as a growth promoter, with increase up to 60% when related to control for all crops. Such results corroborate those reported in our study.

According to Baugh and Escobar 2007, *Trichoderma* action as a growth stimulant is complex and performed by interactions with biochemical factors and production of many enzymes and beneficial compounds for plants. In our work, growth promotion characteristics can be observed.

Pedro et al, 2012, evaluating *Trichoderma spp.* ability in promoting bean plants growth and reducing the severity of bean anthracnose, observed that most efficient isolates can provide increases greater than 30% in dry matter production of plants aerial parts.

### 3.2 Biochemical results

Specific activity of β-1.3 glucanase was higher in plants treated with *T. harzianum ΔEpl1* mutant isolate. In leaves and roots, β-1.3 glucanase activity was estimated as 2.07 Umg^−1^ and 2.57 Umg^−1^ (Table 1), respectively, when inoculated with *S. sclerotiorum*. Without inoculation, leaves and roots presented an average specific activity of 2.27 Umg^−1^ and 2.34 Umg^−1^ (Tables 2 and 3), respectively, which shows a significant result for leaf when compared to other treatments. Regarding roots, differences were observed only in relation to the control treatment, and not in comparison to the wild isolate. Such result was also observed in plants with pathogens, when neither the deletion of *ΔEpl1* or overexpression of aquaporin proteins affected specific activity of β1.3-glucanase. This behavior was also observed for chitinase specific activity, which did not present variation between plants inoculated with mutant and wild isolates.

**Table 1.**
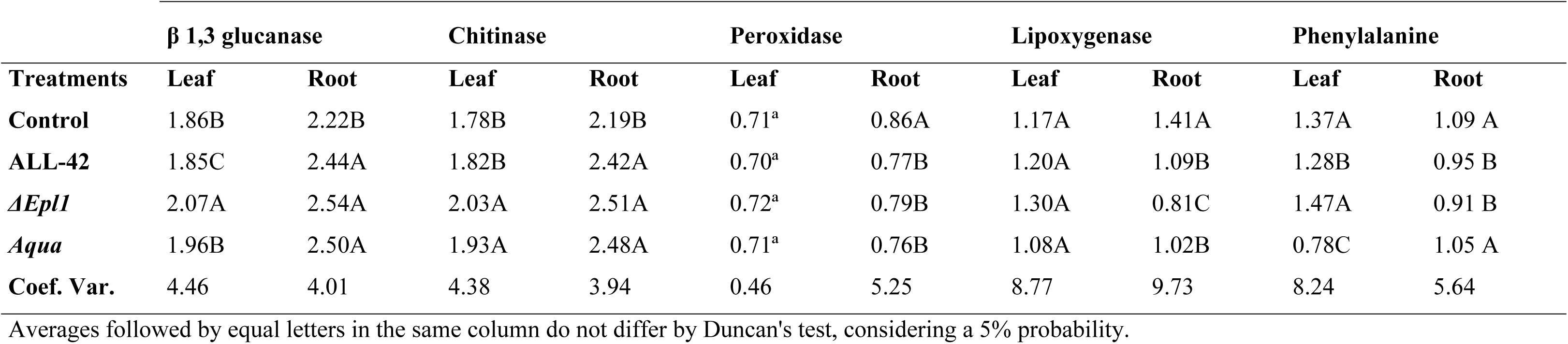
Specific activity (units mg^−1^) of β-1.3-glucanase, chitinase, peroxidase, lipoxygenase and phenylalanine ammonia lyase enzymes, on leaves and roots of common bean plants grown after seed treatments with wild and mutant *Trichoderma harzianum* strains.

**Table 2.**
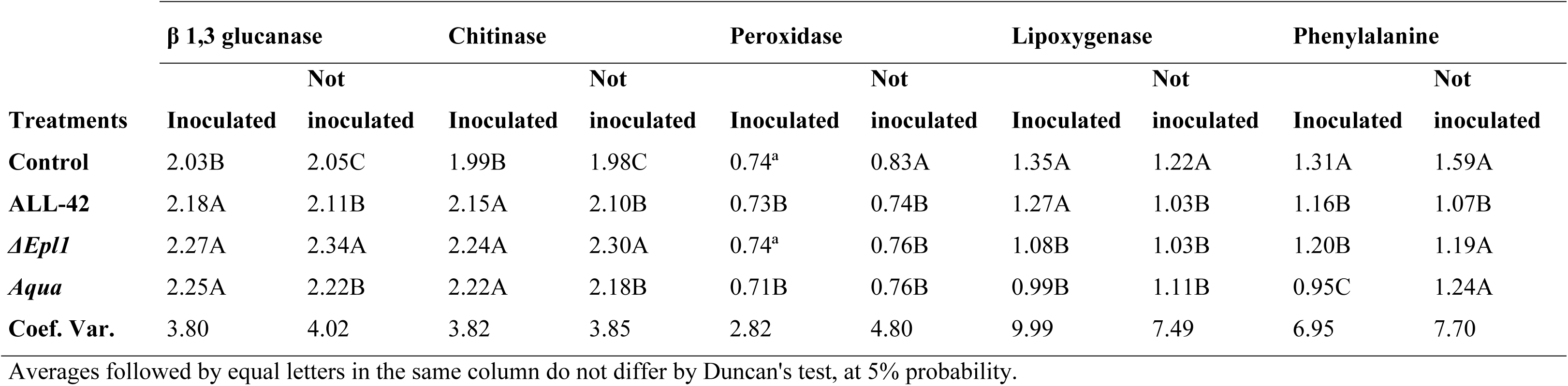
Specific activity (units mg^−1^) of β-1.3-glucanase, chitinase, peroxidase, lipoxygenase and phenylalanine ammonia lyase enzymes in common bean plants grown after seed treatments with wild and mutant *Trichoderma harzianum* strains, inoculated or not with *Sclerotinia sclerotiorum* at the blooming stage.

**Table 3.** Average specific activity (units mg^−1^) of β-1.3-glucanase, chitinase, peroxidase, lipoxygenase and phenylalanine ammonia lyase enzymes on common bean leaves and roots, from plants grown after seed treatments with wild and mutant *Trichoderma harzianum* strains, inoculated or not with *Sclerotinia sclerotiorum* at the blooming stage.

| Treatments | $\beta$ 1.3 glucanase | | Chitinase | | Peroxidase | | Lipoxygenase | | Phenylalanine | |
| --- | --- | --- | --- | --- | --- | --- | --- | --- | --- | --- |
|  | Leaf | Root | Leaf | Root | Leaf | Root | Leaf | Root | Leaf | Root |
| Inoculated | 1.92 A | 2.44 A | 1.89 A | 2.41 A | 0.72 A | 0.84 A | 1.12 B | 1.09 A | 1.34 A | 0.97 B |
| Not inoculated | 1.95 A | 2.41 A | 1.88 A | 2.39 A | 0.71 A | 0.75 B | 1.26 A | 1.08 A | 1.30 A | 1.03 A |

Averages followed by equal letters in the same column do not differ by Duncan's test, at 5% probability.

Attempts to explore antagonist fungi, such as *Trichoderma* species with biocontrol potential, have recently led to the proposal that, in addition to its recognized antifungal properties, these organisms can also be used as elicitors of plant defense reactions, leading to expression of products related to plant defense (LORITO et al., 2010).

Glucanase is an important defense enzyme, especially towards oomycetes, since they have cellulose and glucan as the major cell wall components. Our results corroborate with previous reports that showed increased levels of glucanase during *Trichoderma*-mediated induction in many host-pathogen systems. For instance, Vieira (2013) observed increased glucanase activities associated to induced resistance promoted by *Trichoderma* spp. towards *Phytophthora capsici* in pepper, and of *T. roseum* towards *Macrophomina phaseolina* in chickpea.

Plants treated with *Trichoderma harzianum ΔEpl1* and *Aqua* mutant isolate presented no difference for chitinase activity, once leaf results were 2.03 Umg-1 and 1.93 Umg-1 and root results were 2.51 Umg-1 and 2.48 Umg^−1^, respectively (Table 1). Plants infected or not with *S. sclerotiorum* also exhibit no differences between them, except when compared to control, being the results with pathogen (2.24 Umg-1 and 2.22 Umg^−1^) and without pathogen (2.30 Umg^−1^ and 2.18 Umg^−1^) (Table 2 and 3).

Saksirirat et al. 2009, worked with fifteen *Trichoderma spp.* isolates to control *Xanthomonas campestris pv. Vesicatoria*, what showed a 69.32% decrease in symptoms. He observed that four isolates of *T. harzianum* and *T. asperellum* species induced chitinase activity in the leaves, showing that these species are promising for resistance induction in tomato plants. The same authors reported that this enzyme is known to play an important role in degradation of fungus cell walls. In addition, they are linked to pathogenesis-related proteins (PR proteins) in several plants, including beans.

Peroxidase and lipoxygenase enzymatic activities did not present a significant difference in the leaf when compared to other treatments, whereas in the root and in plants with and without pathogens, control presented higher results, once the results are 0.86 Umg^−1^, 0.74 Umg^−1^ and 0.83 Umg^−1^ for peroxidase and 1.41 Umg^−1^, 1.35 Umg^−1^ and 1.22 Umg^−1^ lipoxygenase. This can propose that *Trichoderma* isolate under these conditions has no effect on peroxidase specific activity of the enzyme (table 1, 2 and 3).

Similar results for peroxidase activity were obtained by Silva et al. (2011), who observed the effect of 60 *Trichoderma spp.* isolates in induction of systemic resistance to anthracnose in cucumber plants. They found no significant difference in relation to activation of the enzyme peroxidase in the plant. This work presented lower values for the isolates when compared to control, which evidences suppression of peroxidase enzymatic activity caused by the action of the *Trichoderma* isolates. Corroborating these results, Dildey et al. 2013, observed no activity of peroxidase induction in common bean against the *Macrophomina phaseolina* pathogen, in the inoculation of different *T. harzianum* isolates through seeds.

Dildey. 2014, when studying *Trichoderma*-bean organisms interaction, verified that TM1, TM4 (*T. virens*), TOD2A and TOD2B (*T. longibrachiatum*) isolates resulted in peroxidase enzymatic suppression, results similar to what was found in the present study. It is suggested that such results were due to a survival strategy or defense by the antagonist, because they prevented activation of the peroxidase enzymatic arsenal.

Plants treated with *Trichoderma harzianum ΔEpl1* mutant isolates showed higher activities of the enzyme phenylalanine anion lyase with 1.47 umg^-1^ in the leaf. However, they did not present significant difference when related to control. In root and in plants with and without pathogen inoculation, results presented higher rates of enzymatic activities in control (1.09 Umg^−1^, 1.31 Umg^−1^ and 1.59 Umg^−1^), suggesting that these plants possibly have no induction in the production of phenylalanine ammonia lyase enzyme (Table 1, 2 and 3).

Phenylalanine ammonia lyase is well established as an important plant defense enzyme induced by *Trichoderma* treatment in several host pathogen models. Phenylalanine ammonia lyase was significantly increased during induction of host resistance by *Trichoderma* species against *Rhizoctonia solani* in sunflower and *F. oxysporum* and *A. alternata* in a work conducted by Surekha, 2014. These results do not corroborate with those found in this work.

Jayalakshmi et al. 2009, by testing *T .harzianum* as a defense elicitor in chickpea (*Cicer arietinum L.*), against wilt caused by *Fusarium oxysporum f. sp. ciceri.* could conclude that *T. harzianum* L1 isolate showed an increase in phenylalanine ammonia lyase activity, differently from the results in the present study.

França et al. 2017, when treating bean plants with *Trichoderma spp.* isolates, aiming the resistance induction, observed that among these, *T. harzianum* ICB05 was the most important one, promoting a notorious increase in peroxidase and phenylalanine ammonia-lyase activity. Such results were not found in the present study.

In our study, peroxidase enzyme activity, with 0.84 umg^-1^ in root, showed higher levels in the pathogen presence. Lipoxygenase enzyme activity was higher in leaf without pathogen, with 1.26 umg^-1^ and phenylalanine ammonia lyase enzyme activity in root, with 1.03 Umg^-1^. Other treatments did not result in significant differences in enzymes activity levels, when related to *S. sclerotiorum* presence or absence (Table 3).

Results showed that plants inoculated with *S. sclerotiorum* exhibit an increase in peroxidase activity. Defense-related enzymes, especially peroxidase, include pathogen spread through formation of polymerized phenolic barriers near infection sites and trigger the synthesis of anti-nutritive, antibiotic and cytotoxic compounds, leading to increased resistance against pathogens (LI L E STEFFENS, 2002). Several papers have emphasized the role of peroxidase in the resistance induced by *Trichoderma* in cultivated plants.

Peroxidase and phenylalanine ammonia lyase increased activities corresponded to resistance of *Macrophomina phaseolina* species in peanut, *Rhizoctonia solani* in sunflower and *Fusarium oxysporum f. sp. lycopersici* in tomato plants (SINGH et al, 2014). This feature is due to *Trichoderma harzianum* NBRI- 1055 induction, obtained by the increased production of defense enzymes, including peroxidase and phenylalanine ammonia lyase activities. Similarly, Christopher, 2010 observed that peroxidase activities were significantly higher in tomato plants during resistance induced by *Trichoderma virens* against *Fusarium* wilt, caused by *Fusarium oxysporum f. sp. lycopersicy*.

Mechanisms used by Trichoderma species to elicit plant defense responses are not completely understood; however, studies on differential gene expression and changes in protein expression patterns of leaves and roots of host plants during association with *Trichoderma* species were performed (DJONOVI et al., 2006). Synthesis of PR protein, involving peroxidases, chitinases, β-1.3-glucanases and lipoxygenase, can be considered as one of the most evident alterations in plant-pathogen interaction. Other side infection responses include increased expression of genes related to phytoalexins and phenylalanine ammonia lyase synthesis, within lignin deposition and higher salicylic acid (SA) levels (VERBERNE et al, 2000).

When plants are exposed to pathogens, along with biochemical and molecular response, plant tissues mechanical strengthening may contribute to limit pathogen spread (Chowdhury et al., 2014; Rao et al., 2015). Plant defense response and resistance against fungal pathogens was found to partially come from cell wall strength stimulation by lignin and callus formation in plants (Nikraftar et al., 2013). Simultaneous participation of peroxidase, chitinase and lignin formation in tomato and rice protection against *R. solani* was described by Taheri and Tarighi, 2012.

### Principal component analysis

The three main components of PCA were responsible 45.74%, 26.31% and 11.83% of the total variance, respectively, accounting on 83.88% of data variation (Figure 2). In this joint analysis, there were slight differences between inoculated and non inoculated plants, regarding the control and *ΔEpl1* treatments. In contrast, the ALL-42 wild strain and its *Aqua* mutant showed distinct responses to inoculation with *S. sclerotiorum*. Leaf area, root length, root volume and phenylalanine ammonia lyase and lipoxygenase enzymes in the roots are responsible for a large variation in the data. Plant growth variants, as leaf area, root length and volume were negatively correlated with lipoxygenase and phenylalanine ammonia lyase enzymatic activity in leaf, chitinase and □1.3 glucanase in leaf and root. These correlations indicate that plant growth promotion is inversely proportional to enzymatic activity of these enzymes..

**Figure 2:**
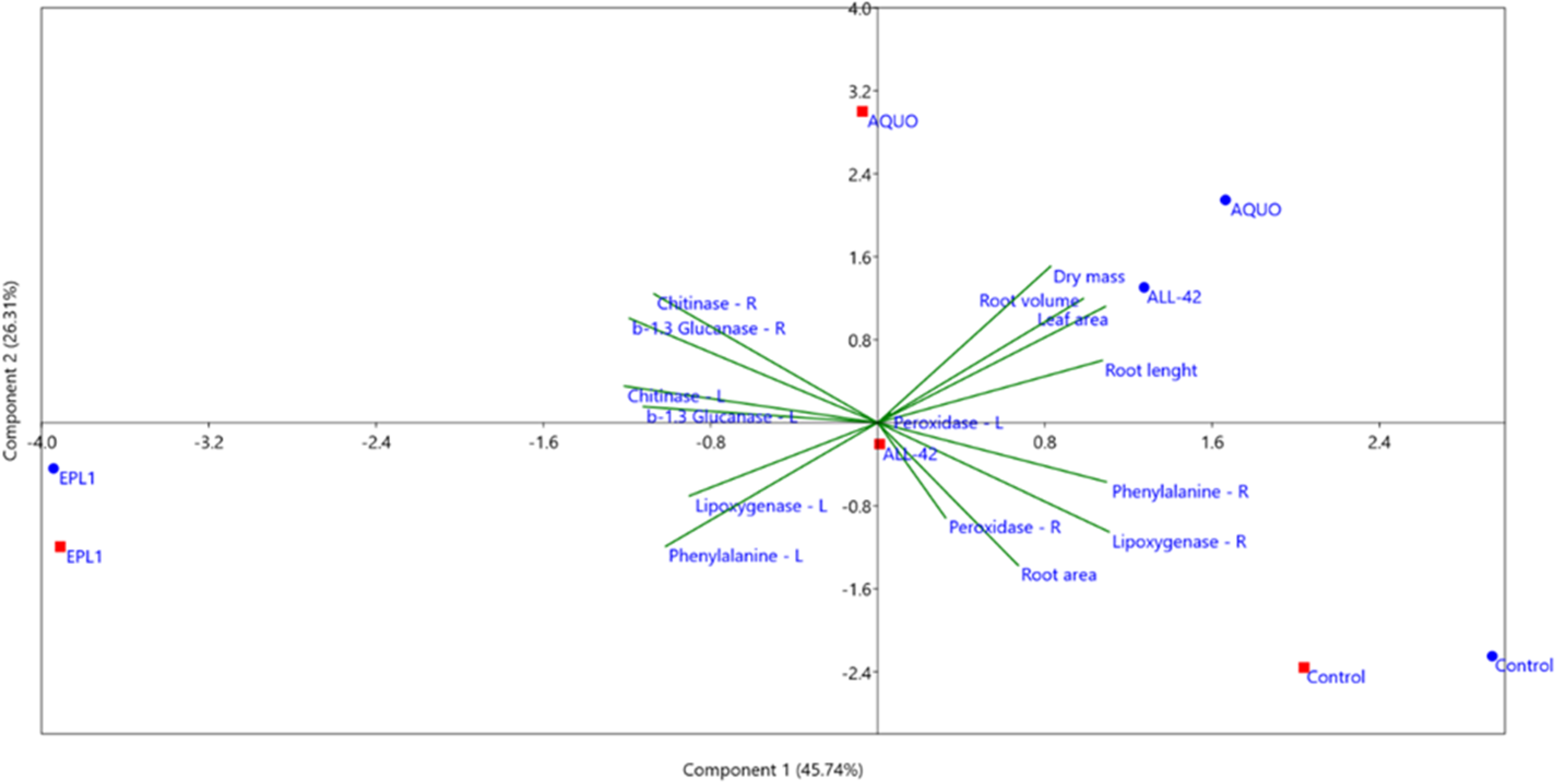
Principal component analysis biplot with relationships between physiological traits (leaf area, dry mass, root volume, root length and root surface) and enzymes related to resistance induction to *Sclerotinia sclerotiorum*, on common bean plants grown after seed treatments with wild and mutant *Trichoderma harzianum* strains. Red squares indicate plants inoculated with *S. sclerotiorum*, while blue circles regard to non-inoculated plants.

## 4. Conclusions

As for plant and *Trichoderma* interaction, *Sclerotinia sclerotiorum* pathogen showed no influence on plant growth or resistance induction. *Trichoderma harzianum Aqua* mutant isolate promotes common bean plant growth, in comparison to its wild strain ALL-42. *Trichoderma* mutant isolates do not induce resistance of common bean plants to *S. sclerotiorum*.

## Acknowledgments

RSB was were funded by Research Foundation of the State of Goiás (FAPEG, GO, Brazil). The authors wish to thank Dr. Robert Pogue by the English revision of the text and for fruitful discussions.

